# Small, but mitey: Investigating the molecular genetic basis for mite domatia development and intraspecific variation in *Vitis riparia* using transcriptomics

**DOI:** 10.1101/2024.03.04.583436

**Authors:** Eleanore J. Ritter, Carolyn D. K. Graham, Chad Niederhuth, Marjorie Gail Weber

## Abstract

• Here, we investigated the molecular genetic basis of mite domatia, structures on the underside of leaves that house mutualistic mites, and intraspecific variation in domatia size in *Vitis riparia* (riverbank grape).

• Domatia and leaf traits were measured, and the transcriptomes of mite domatia from two genotypes of *V. riparia* with distinct domatia sizes were sequenced to investigate the molecular genetic pathways that regulate domatia development and intraspecific variation in domatia traits.

• Key trichome regulators as well as auxin and jasmonic acid are involved in domatia development. Genes involved in cell wall biosynthesis, biotic interactions, and molecule transport/metabolism are upregulated in domatia, consistent with their role in domatia development and function.

• This work is one of the first to date that provides insight into the molecular genetic bases of mite domatia. We identified key genetic pathways involved in domatia development and function, and uncovered unexpected pathways that provide an avenue for future investigation. We also found that intraspecific variation in domatia size in *V. riparia* seems to be driven by differences in overall leaf development between genotypes.

## INTRODUCTION

Mutualisms between plants and arthropods have evolved repeatedly across evolutionary time (Blattner *et al*., 2001; Bronstein *et al*., 2006), promoting the evolution of unique, heritable structures in plants that attract, reward, or protect mutualists (Romero & Benson, 2005; Bronstein *et al*., 2006). Investigating the genetic basis of mutualistic structures provides a valuable lens for understanding how mutualisms evolved. Mite domatia (hereafter “domatia”) are tiny mutualistic plant structures on the underside of leaves that provide shelter for beneficial mites that have received relatively little attention from the genetic perspective despite being produced by many woody plant species. Domatia facilitate a bodyguard mutualism between plants and mites: mites benefit from the refuge provided by the domatia, which protects them from predators (Grostal & O’Dowd, 1994; Norton *et al*., 2001; Faraji *et al*., 2002a,b; Romero & Benson, 2005), and in return plants receive protection from pathogenic fungi and/or herbivory via fungivorous and/or predacious mites (Agrawal & Karban, 1997; Norton *et al*., 2000; Romero & Benson, 2004). Domatia are common defenses in natural systems: they are present in over 5,000 plant species and make up a large proportion of woody plant species in temperate deciduous forests (e.g., ∼50% of woody plant species in forests in Korea (O’Dowd & Pemberton, 1998) and Eastern North America (Willson, 1991)). They are present in several crop plants and have been studied as a pest control strategy in agriculture (Romero & Benson, 2005; Barba *et al*., 2019). Yet, despite their agricultural and ecological importance, we know relatively little about the genetic underpinnings of mite domatia in plants.

The genus *Vitis* is a powerful group for studying the genetics of domatia due to the heritable variation of domatia presence and size (English-Loeb *et al*., 2002; Graham *et al*., 2023) and the genetic and germplasm resources available. In *Vitis,* domatia are constitutive, small, dense tufts of trichomes covering a depression in the leaf surface in the abaxial vein axils, termed “tuft” domatia. Norton *et al*. (2000) demonstrated that domatia in *Vitis riparia,* a wild grapevine species with relatively large domatia, led to a 48% reduction in powdery mildew in comparison to *V. riparia* plants with blocked domatia, which were inaccessible to mites. Given how effective domatia are as biological control agents in this system, there is interest in understanding domatia in domesticated grapevine (*Vitis vinifera*) and related species.

Two previous studies investigated the genetic basis of domatia in *Vitis* (Barba *et al*., 2019; LaPlante *et al*., 2021). Barba *et al*. (2019) measured mite abundance, domatia, and general trichome traits in the segregating F_1_ family of a complex *Vitis* hybrid cross. They identified multiple QTLs influencing domatia-related traits, including a major QTL on chromosome 1. They also found additional support for a relationship between overall leaf and leaf trichome development, previously demonstrated in *Vitis* (Chitwood *et al*., 2014). LaPlante *et al*. (2021) investigated the genetic basis of trichome and domatia traits in a genome-wide association study (GWAS) using a common garden of *V. vinifera* cultivars. They identified a single nucleotide polymorphism (SNP) associated with domatia density near several candidate genes on chromosome 5. Only one gene was identified that was shared in both studies: *Glabrous Inflorescence Stems 2* (VIT_205s0077g01390), which is thought to encode a zinc finger protein that regulates trichome development (LaPlante *et al*. 2021). The minimal overlap between the two studies is likely due to differences in the scale of genetic diversity investigated in QTL mapping and GWAS. As a result, the various molecular pathways involved in domatia development remain relatively unknown.

While little is known about the development of tuft domatia specifically, work in related structures in other species may provide clues regarding the genes involved in domatia development. Substantial work has characterized the genes involved in the development of trichomes, which are an essential component of tuft domatia. The molecular pathways involved in trichome development have yet to be elucidated in *Vitis*, though they have been characterized in other angiosperms. In addition, previous work has characterized the genes involved in another form of domatia that house ants, called tuber domatia. Tuber domatia are functionally like tuft domatia in providing shelter for mutualistic arthropods in return for defense, but are tubers formed from stem tissue. The genes involved in domatia development may overlap with those previously implicated in trichome and tuber domatia development, providing additional hypotheses to investigate regarding domatia development in *V. riparia*.

Here, we investigate the molecular genetic mechanisms of development and intraspecific variation in domatia of *V. riparia,* the riverbank grape. We hypothesized that genes differentially expressed in *V. riparia* domatia (i) share similarities with pathways previously identified in trichome development, including TFs and cell wall modification pathways (Dong *et al*., 2022; Han *et al*., 2022), (ii) are involved in responses to biotic organisms as has been previously identified with functionally similar tuber domatia (Pu *et al*., 2021), and (iii) involve auxin-signaling due to its role in both trichome (Han *et al*., 2022) and tuber domatia development (Pu *et al*., 2021). We also hypothesized that intraspecific variation in domatia size in *V. riparia* may be driven by differences in overall leaf morphology, as previous work has demonstrated a link between leaf morphology and trichomes in *Vitis* (Chitwood *et al*., 2014; Barba *et al*., 2019). We sequenced the transcriptomes of domatia in two *V. riparia* genotypes that differ in their investment in domatia alongside control leaf tissue to identify key pathways involved in domatium development in *Vitis* and landmarked leaves from these two genotypes and compared leaf shapes to identify possible morphological differences that may impact domatia traits.

## MATERIALS AND METHODS

### Plant material

We pre-selected two genotypes of *V. riparia* that were identified in a previous study to nearly span domatia size variation in the species (English-Loeb & Norton, 2006): genotype 588711, a large domatia genotype (hereafter LDG) and genotype 588710, a small domatia genotype (hereafter SDG). Hardwood cuttings of each genotype were sourced from the United States Department of Agriculture - Germplasm Resource Information Network (USDA-GRIN) repository in February of 2022. The genotypes were initially collected in Wyoming, USA (SDG) and Manitoba, Canada (LDG). Budding cuttings were potted in March of 2022 and grown in a common garden greenhouse at Michigan State University in East Lansing, Michigan, USA. The vines were watered daily during the growing season with roughly 200 ppm Peters Excel pHLow 15-7-25 High-Mag/High K with Black Iron fertilizer (ICL, Tel-Aviv, Israel) dissolved in water. No pesticides were applied to the plants.

### Characterizing domatia traits

To confirm that the two domatia genotypes used in our study statistically differed in domatia size, we collected, dried, and pressed leaves and scored domatia traits using a dissection microscope. Domatia traits were measured on 1-3 fully expanded leaves per plant for five replicate plants per genotype (25 leaves total). To evaluate how domatia size and density changed throughout leaf ontogeny, we collected 5-7 leaves from five plants of each genotype selected to represent the entire leaf lifespan from bud burst to full expansion (55 leaves total). The leaves were scanned while fresh using a CanoScan 9000F Mark II (Canon U.S.A., Inc.) at 1200 DPI.

Two aspects of domatia were measured for both datasets: hair density and size. Domatia hair density scores were assigned using a nine-point scale where 0 represents no hair, and 9 represents a densely packed domatium with no leaf surface visible underneath (Graham *et al*., 2023). This scale was adapted from the OIV code O-085/U-33 scale, a standard scale for measuring leaf hair density used by grape breeders (IPGRI *et al*., 1997). Domatium radius was used as a proxy for domatia size, measured on pressed leaves using an ocular micrometer, and in pixels and converted to mm using the software ImageJ 1.54d on leaf scans (Schneider *et al*., 2012).

A Mann-Whitney test and a Welch’s two-sided t-test were used to test for differences in domatia density and size, respectively, between the two genotypes. The phenotypic data were plotted using ggplot2 v3.4.2 (Wickham, 2016) and cowplot v1.1.1 (Wilke, 2021) in R. The R package ggsignif v0.6.4 was used to add significance bars (Constantin & Patil, 2021). We evaluated domatia development throughout leaf expansion and identified our RNA sampling timepoints by comparing domatia size on leaves that had not fully expanded (ranging from 2.6-5.8 cm in width) to domatia size on larger leaves using a Welch’s two-sided t-test (Supporting Information Fig. S1). All R analyses (including downstream analyses) were run using R v4.2.2 (R Core Team, 2022) and RStudio v2022.12.0.353 (RStudio Team, 2022).

### Tissue collection, RNA extraction, and sequencing

RNA samples were collected across two consecutive days at the same time (within one hour from start to finish) from the same plants used for characterizing domatia traits. Samples were collected from young leaves that had not fully expanded (ranging from 2.6-5.8 cm in width), as our domatium ontogeny data demonstrated that domatia were still developing during this time (*P* < 0.001, Supporting Information Fig. S1). The samples were collected using a circular 1.5 mm hole puncher on domatia and control tissues. Domatia samples were collected at the center of domatia, avoiding veins, and control samples were on laminar tissue 1.5 mm away from the domatia (Supporting Information Fig. S2). Domatia and control samples were taken from the same leaves, with the first sample (domatia or control) alternated. Between 20-23 tissue samples were taken per plant (across 2-3 leaves) for both domatia and control. Samples were immediately frozen in liquid nitrogen and stored at -80℃.

Samples were pooled into grinding tubes by plant and tissue type and ground using two metal balls per tube in a SPEX™ SamplePrep 2010 Geno/Grinder 2010 at 1750 strokes/minute for 30 seconds. RNA was immediately extracted from pooled samples after grinding using a Spectrum™ Plant Total RNA Kit from Sigma-Aldrich (STRN250) and an On-Column DNase I Digestion Set from Sigma-Aldrich (DNASE70-1SET) to remove DNA. Concentrations were checked with a Qubit High Sensitivity (HS) RNA Assay Kit (Q32852) and an Invitrogen Qubit 4 Fluorometer. The quality of the samples was analyzed using an Agilent 4200 TapeStation, and samples with low quantity and/or quality were discarded.

Fourteen RNA samples with sufficient quantity and quality - paired domatia-control samples from three 588710 plants and four 588711 plants - were prepared for RNA-sequencing (RNA-seq) using the Illumina Stranded mRNA Library Preparation, Ligation Kit with IDT and Illumina Unique Dual Index adapters following the manufacturer’s recommendations, except half volume reactions were used. The quantity and quality of the prepared libraries were checked using a Qubit HS dsDNA Assay Kit (Q32851) and the High Sensitivity D1000 ScreenTape assay, respectively. The libraries were pooled and sequenced using one Illumina S4 flow cell lane in a 2x150bp paired-end format and a NovaSeq v1.5 reagent kit (300 cycles). Base calling was done by Illumina Real Time Analysis (RTA) v3.4.4. The output of RTA was demultiplexed and converted to FastQ format with Illumina Bcl2fastq v2.20.0.

### RNA read processing and mapping

FastQC v0.11.9 was used to check the quality of the raw RNA-seq reads (Andrews, 2010). One sample failed poly-A capture during library preparation, so both samples from the failed library were excluded from downstream analysis, leaving three paired RNA-seq samples for each genotype. RNA-seq reads were trimmed using Trimmomatic v0.39 with the *-phred33* flag and the provided NexteraPE-PE.fa adapter file (Bolger *et al*., 2014). The quality of the trimmed reads was confirmed using FastQC v0.11.9 (Andrews, 2010). After read trimming, there were 26-51 million reads per sample (Supporting Information Table S1).

Trimmed RNA-seq reads were mapped to the *V. riparia* genome (Girollet *et al*., 2019) using STAR v2.7.0c (Dobin *et al*., 2013). STAR genome index files were generated by running STAR with the *V. riparia* genome and annotations, using the flags *--runMode genomeGenerate* and *--genomeSAindexNbases 13*. The trimmed reads were mapped using STAR with the generated index files and the *--quantMode GeneCounts* flag. Between 23-46 million reads (84.9-91.6% of trimmed reads) mapped to the *V. riparia* genome (Supporting Information Table S1).

### Differential expression analysis

Mapped read count outputs from STAR were used for differential expression analysis using DESeq2 v1.38.3 (Love *et al*., 2014) downloaded from Bioconductor (Huber *et al*., 2015). DESeq2 was run using the *design = ∼genotype + tissue + genotype:tissue*, where tissue was either control or domatia. Genes were considered differentially expressed between groups if the absolute value log2 fold change was greater than 1 and the adjusted *P*-value was less than 0.05 (Supporting Information Tables S2-S5). Heatmaps displaying differentially expressed genes (DEGs) were created using the ComplexHeatmap v2.14.0 package (Gu, 2022).

### GO term enrichment analysis

Gene ontology (GO) term enrichment analysis was performed on upregulated DEGs. GO term annotations are more robust in Arabidopsis (*Arabidopsis thaliana*), so we utilized Arabidopsis orthologs for our GO term enrichment analysis. We identified Arabidopsis orthologs to *V. riparia* genes using protein sequences for both species with diamond v2.0.15.153 (Buchfink *et al*., 2015) and the following flags: *--iterate, --max-target-seqs 1, and --unal 0*. GO terms of Arabidopsis orthologs for DEGs were used from The Arabidopsis Information Resource (TAIR) (Berardini *et al*., 2015). Interproscan v5.61-93.0 (Jones *et al*., 2014) was used to identify PFAM GO terms based on *V. riparia* protein sequences. GO terms for the Arabidopsis orthologs and those generated by PFAM were concatenated, and parent GO terms were added using GO files from the GO knowledgebase v2023-11-15 (Ashburner *et al*., 2000; Gene Ontology Consortium, 2023). GO term enrichment for upregulated DEGs was assessed by performing a Fisher’s exact test with Benjamini-Hochberg correction using TopGO v2.44.0 (Alexa & Rahnenfuhrer, 2023) from Bioconductor (Huber *et al*., 2015) against the background of *V. riparia* genes. GO terms were considered enriched with a *P*-value < 0.05. GO term enrichment results were plotted using ggplot2 v3.4.2 (Wickham, 2016).

### Leaf landmarking and leaf shape analysis

To test whether differences in leaf shape may be related to differences in domatia size between SDG and LDG, we compared leaf shapes between the two. To do so, leaves used for measuring domatia ontogeny were landmarked manually by placing 21 landmarks described in Bryson *et al*. (2020) on leaf scans using ImageJ v1.53k (Schneider *et al*., 2012). Landmarks were saved as x- and y-coordinates in centimeters. Comparing differences in leaf shape was performed as described in Ritter *et al*. (2023) using the shapes package v1.2.7 (Dryden & Mardia, 2016). A Hotelling’s *T^2^* test was used to test for mean shape differences. Average leaves for each sample and PC values were plotted using ggplot2 v3.4.2 (Wickham, 2016) and cowplot v1.1.1 (Wilke, 2021).

## RESULTS

### Genotypes differ in domatia investment

We characterized domatia traits in two *V. riparia* genotypes previously reported to have different domatia sizes (English-Loeb & Norton, 2006). The two genotypes differed in domatia size, with LDG having larger domatia than SDG (*P* < 0.001) (Fig. 1). The genotypes had similarly dense domatia (*P* = 0.382).

**Fig. 1.**
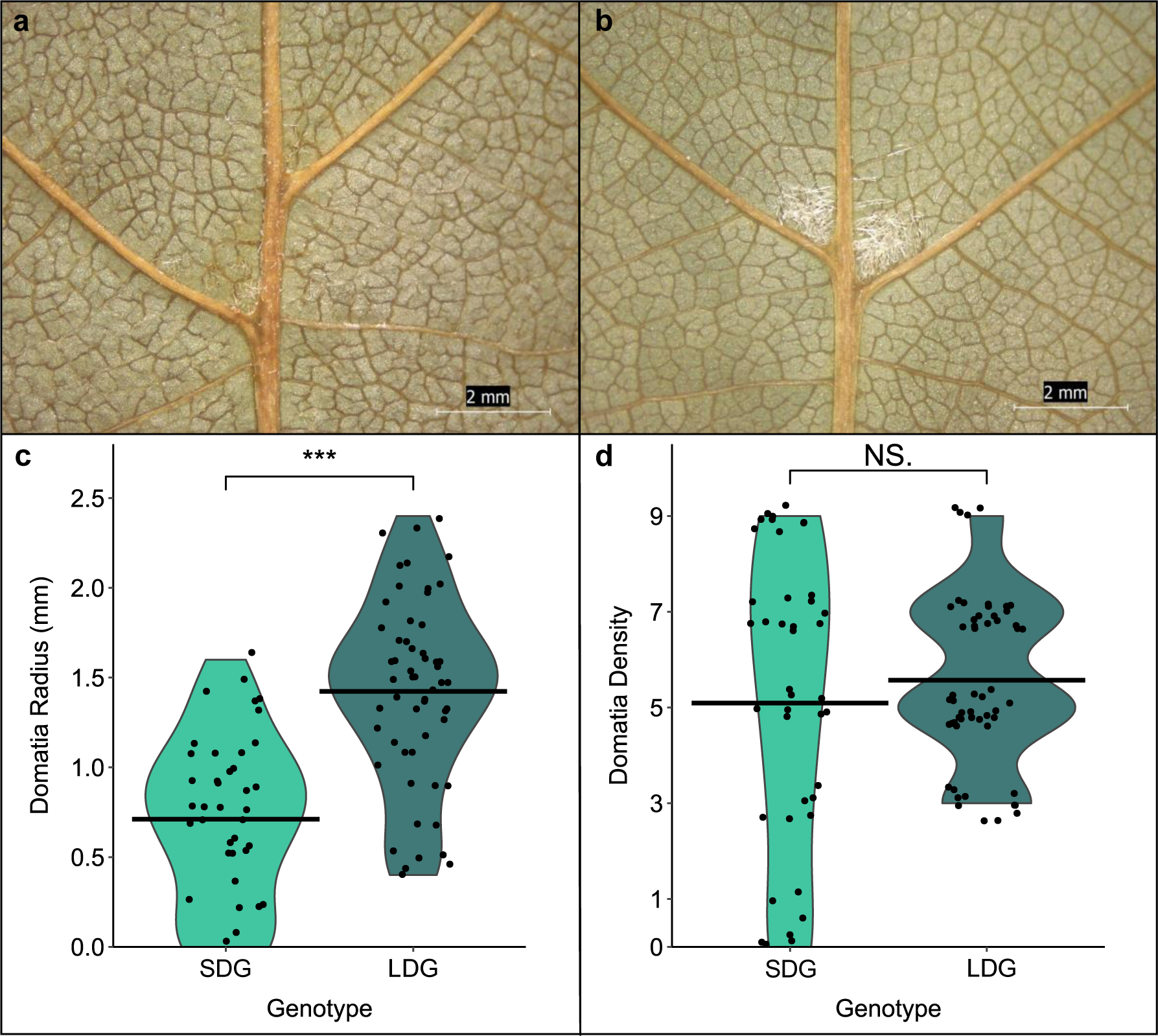
Domatia size and density of *V. riparia* SDG and LDG plants. **(a)** Domatia from the SDG. **(b)** Domatia from the LDG. **(c)** The radius of domatia (mm) from SDG and LDG plants, with the average radius of domatia represented by a black line (\*\*\**P* < 0.001). **(d)** The domatia density of both genotypes, from 0 (representing essentially no domatium present) to 9 (representing a very dense domatium), with the average density for each genotype represented by a black line (NS., *P* = 0.382).

However, the range of domatia density in the SDG (0-9) was greater than the range of domatia density in LDG (3-9) due to both absent and very sparse domatia in the SDG (scored 0 and 1, respectively) (Fig. 1).

### Differentially expressed genes in domatia

Differential expression analysis revealed 1,447 and 759 DEGs in the SDG and the LDG domatia compared to control tissue, respectively. Most DEGs were upregulated in the domatia (88.6% in SDG and 94.9% in LDG). There was substantial overlap of DEGs between the two genotypes, with 538 genes (∼37% SDG, ∼71% LDG) overlapping (Supporting Information Fig. S3).

GO term enrichment analysis revealed 98 and 58 Biological Process (BP) GO terms enriched in SDG and LDG, respectively (Supporting Information Tables S6 and S7). Of these, 39 were shared between genotypes (Fig. 2) and primarily fell into three categories - development, hormone signaling, and responses to stimuli.

**Fig. 2.**
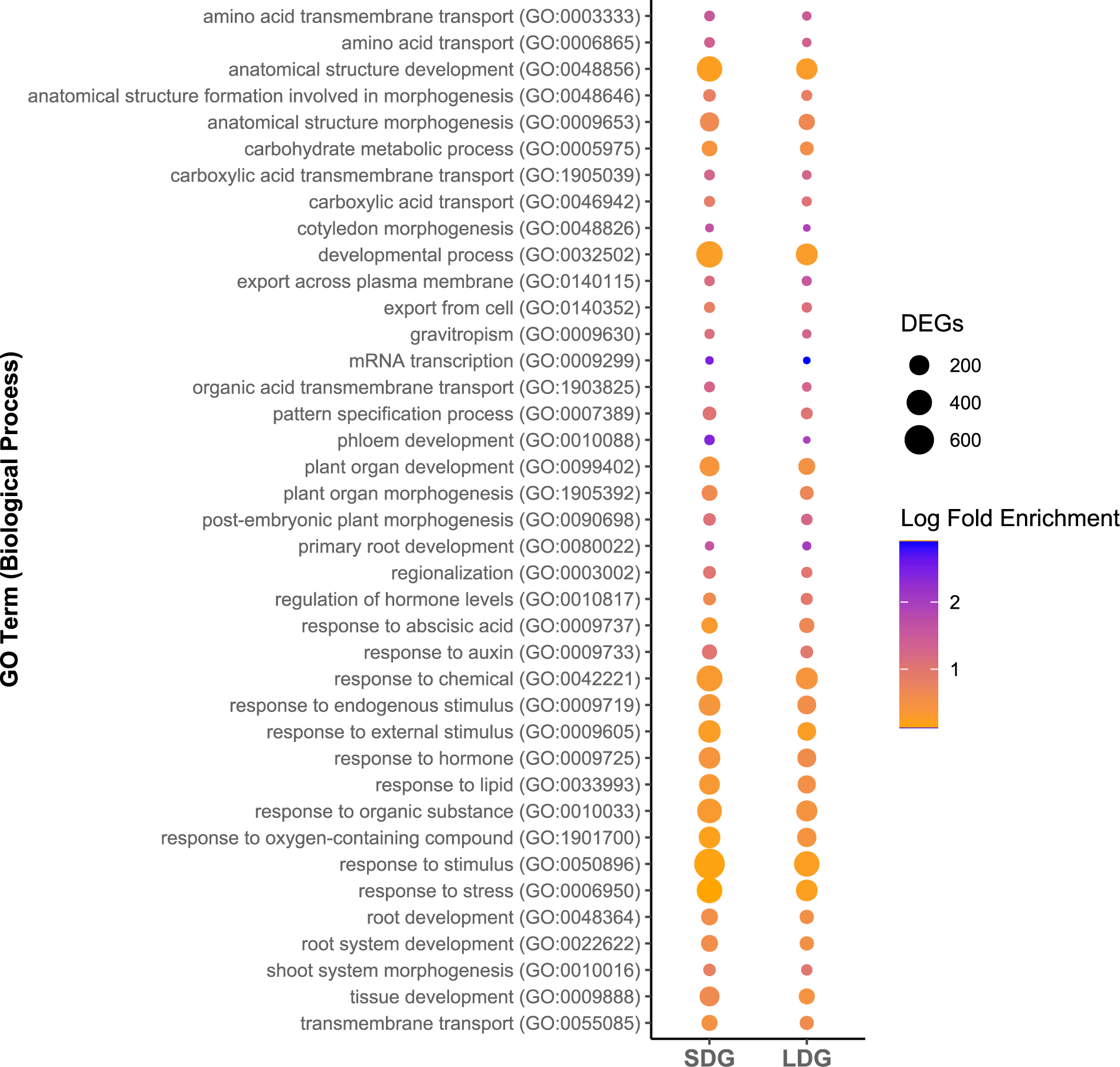
Biological Process gene ontology (GO) terms enriched (*P* < 0.05) in both genotypes in differentially expressed genes (DEGs) upregulated in domatia tissue.

DEGs were also enriched for 7 and 8 Cellular Component (CC) GO terms for SDG and LDG, respectively, with 7 of these shared between the genotypes (Supporting Information Fig. S4), including components in the cell wall, plasma membrane, and regions outside the cell. A total of 23 and 24 Molecular Function (MF) GO terms were enriched for upregulated genes in SDG and LDG, respectively. However, only 12 MF GO terms were enriched in both genotypes (Supporting Information Fig. S5). While the MF GO terms enriched were diverse in function, in general, transporter activity and nucleic acid binding were commonly enriched.

### Regulators of trichome development are upregulated in domatia

We identified multiple TFs upregulated in domatia that have been shown to regulate trichome initiation, including C2H2 ZFPs and SQUAMOSA promoter-binding protein-like (SPLs) (Zhou *et al*., 2013; Yue *et al*., 2018; Zhang *et al*., 2020; Han *et al*., 2022). Five genes encoding C2H2 ZFPs, orthologous to *AtNTT* (two *V. riparia* genes), *AtZFP1*, *AtZFP4,* and *AtZFP6,* were upregulated in domatia in both *V. riparia* genotypes. We also found that the *V. riparia* genes orthologous to *AtSPL13A* and *AtSPL12* are upregulated specifically in domatia of one genotype for SDG and LDG, respectively (Supporting Information Fig. S6).

### Cell wall gene expression is upregulated in domatia

As domatia formation requires both the development of trichomes and the depression in the leaf lamina, we hypothesized that cell wall modification genes would be upregulated in domatia. We found that approximately 7% of genes upregulated in domatia are involved in biosynthetic pathways for cell wall components, predominantly hemicelluloses (namely xylan and xyloglucan), pectin, and lignin (Fig. 3), with upregulated genes in SDG domatia enriched for the biosynthetic processes of all three (Supporting Information Table S6).

**Fig. 3.**
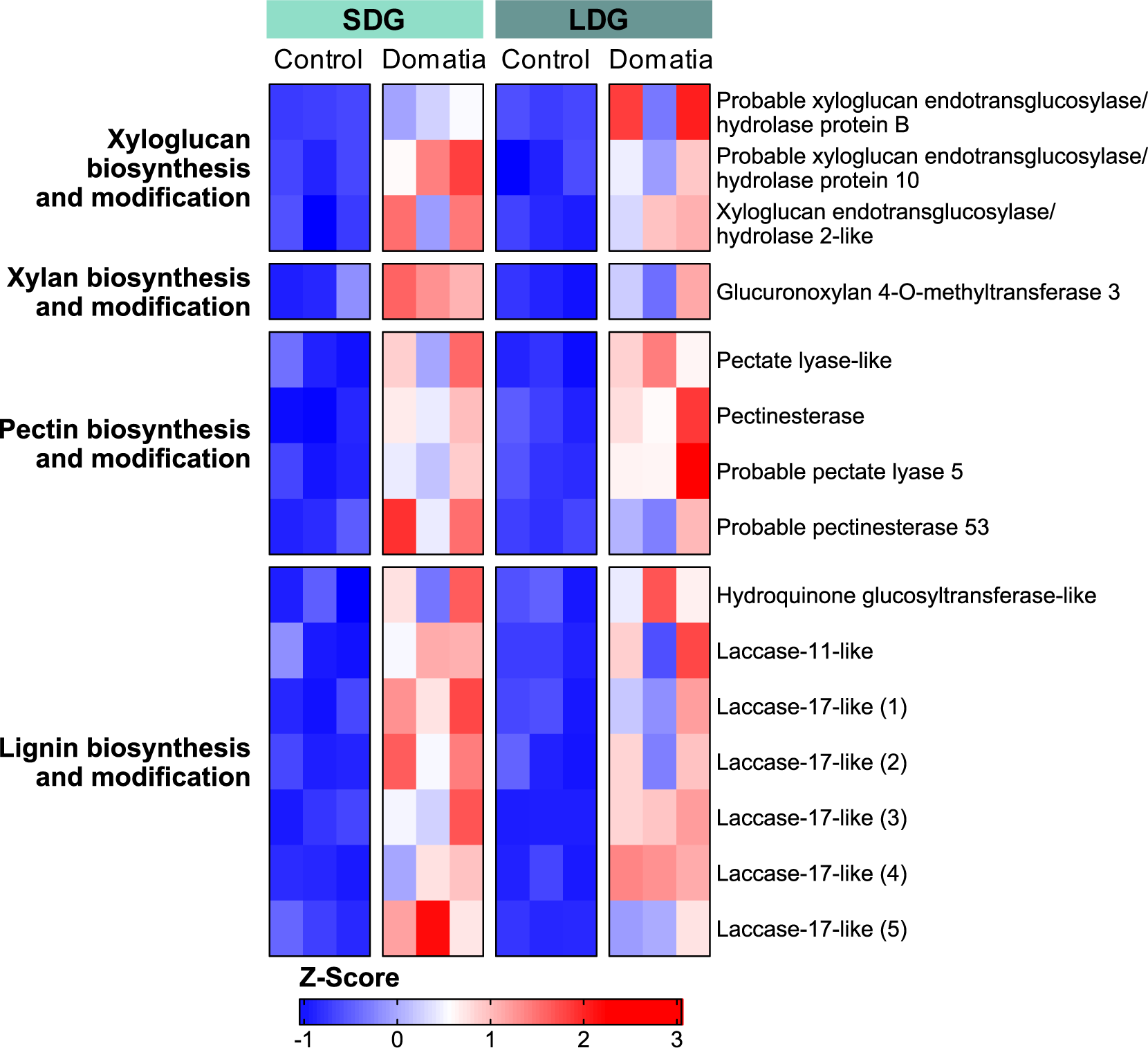
A heatmap of genes upregulated in domatia of both genotypes that are involved in cell wall biosynthesis and modification. Each column represents a biological replicate. The cells are colored by Z-score, with blue representing lower expression and red representing higher expression.

These include many genes that do not overlap between genotypes but are within the same gene families, including two different *V. riparia* orthologs to *AtPARVUS,* a key gene in xylan biosynthesis (Lee *et al*., 2007).

### Upregulated genes in domatia mediate interactions with biotic organisms

As identified in tuber domatium (Pu *et al*., 2021), many genes upregulated in mite domatia are involved in direct defense responses against pathogens. Seventeen genes in SDG and twelve genes in LDG upregulated in domatia are from the *NBS-LRR* family, which are generally involved in pathogen detection (DeYoung & Innes, 2006), with six genes shared between genotypes (Fig. 4).

**Fig. 4.**
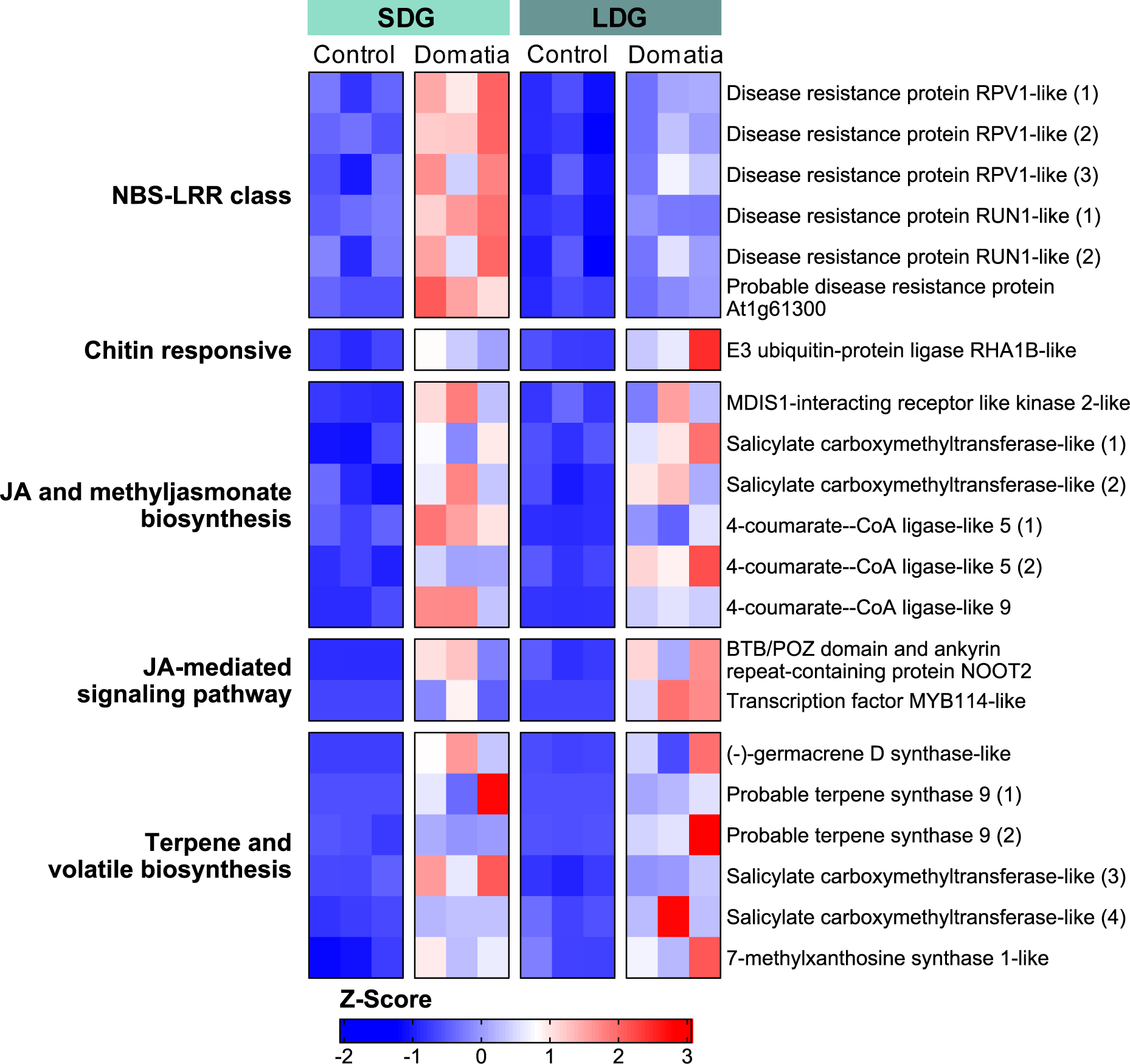
A heatmap of genes upregulated in domatia of both genotypes that are involved in interactions with biotic organisms. Each column represents a biological replicate. The cells are colored by Z-score, with blue representing lower expression and red representing higher expression.

Beyond *NBS-LRR* genes, additional genes involved in pathogen and chitin sensing are upregulated in domatia, including genes encoding E3 ubiquitin-protein ligase RHA1B-like (Fig. 4) and protein LYK5-like (in SDG domatia only).

The jasmonic acid (JA) signaling pathway was previously implicated in both tuber domatia (Pu *et al*., 2021) and trichome development (Han *et al*., 2022) and is upregulated in *V. riparia* domatia for both genotypes. Orthologs to *JA carboxyl methyltransferase* (*JMT*) in Arabidopsis (annotated as encoding salicylate carboxymethyltransferase-like), which plays a crucial role in JA signaling by catalyzing the formation of methyl jasmonate from JA (Seo *et al*., 2001), are upregulated in domatia, with SDG domatia having two and LDG domatia having three *JMT* orthologs upregulated. Other genes thought to be involved in JA biosynthesis are upregulated in domatia as well, including two genes encoding 4-coumarate--CoA ligase-like 5 proteins (orthologous to *AtOPCL1*) and one encoding 4-coumarate--CoA ligase-like 9 (orthologous to AT5G63380) (Fig. 4).

Possibly because of JA signaling, genes involved in terpene and volatile synthesis are upregulated in domatia, including genes shown to mediate plant-arthropod interactions. These include the orthologs to *TPS03*, *TPS21*, and *TPS24* in Arabidopsis, all of which are involved in the synthesis of volatile compounds. In addition, two genes upregulated in domatia that encode Salicylate carboxymethyltransferase-like (3-4) are orthologous to the Arabidopsis gene encoding AtBSMT1, which is involved in the production of the volatile compound Methyl Salicylate (MeSa) (Fig. 4).

### Domatia development is likely regulated by auxin signaling

Auxin signaling has been implicated in regulating both trichome development (Han *et al*., 2022) and tuber domatia development (Pu *et al*., 2021) and seems to play a role in domatia in *V. riparia,* which have genes upregulated at multiple steps in the auxin signaling pathway (Fig. 5).

**Fig. 5.**
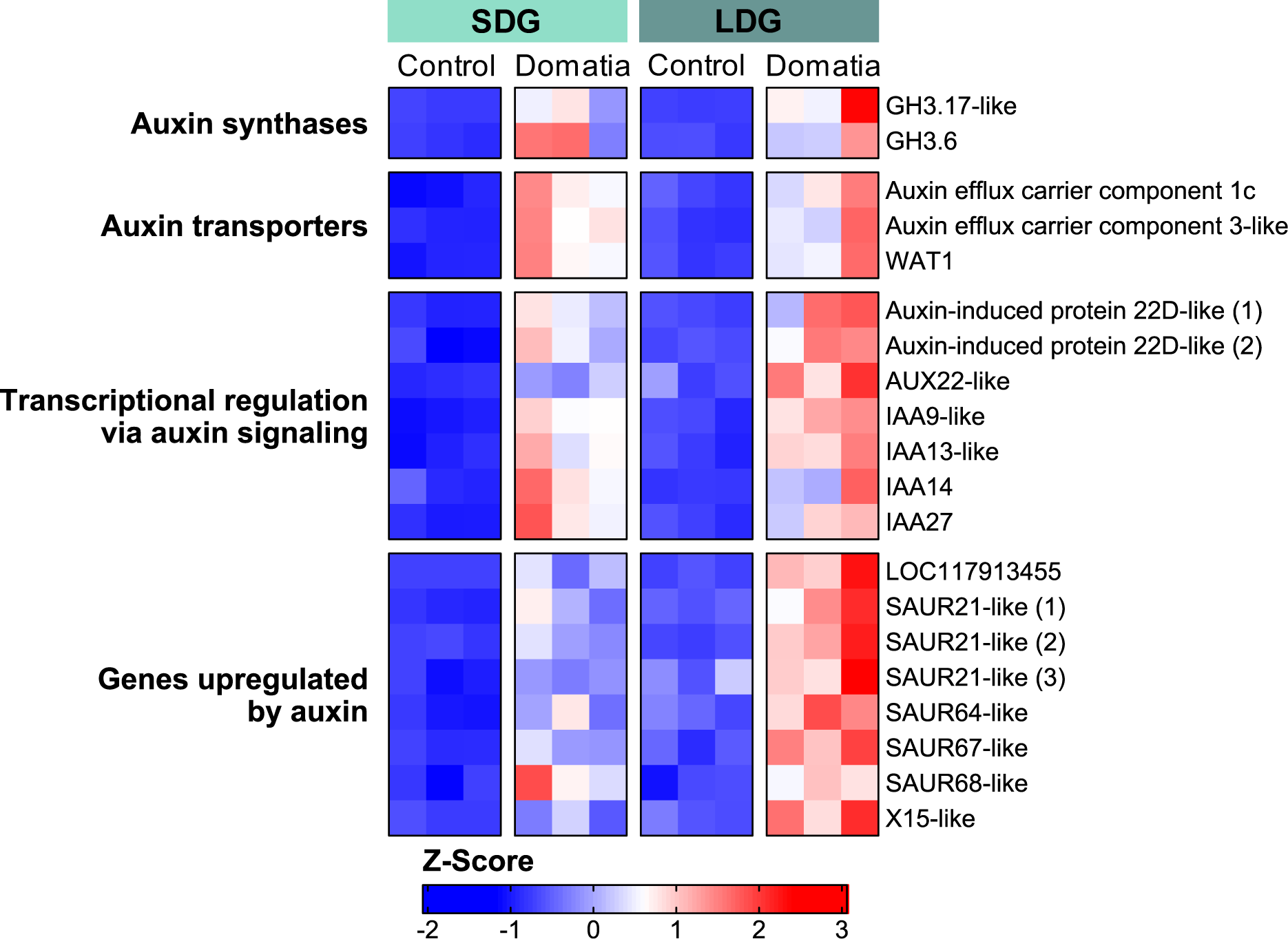
A heatmap of genes upregulated in domatia of both genotypes that are involved in auxin signaling. Each column represents a biological replicate. The cells are colored by Z-score, with blue representing lower expression and red representing higher expression.

Two genes encoding auxin synthases (GH3.6 and GH3.17) are upregulated, suggesting that auxin is actively produced during domatia development. In addition, three auxin transporters are upregulated in domatia in both genotypes, including genes encoding auxin efflux carrier components. Several genes involved in auxin responses downstream are upregulated in both genotypes as well, including six *auxin/indole-3-acetic acid* (*Aux/IAA*) genes, which are both involved in transcriptional regulation via auxin signaling (Ulmasov *et al*., 1997; Leyser, 2018), as well as two genes encoding auxin response factors (ARFs). Genes typically upregulated by auxin are also upregulated in domatia of both genotypes, including eight *small auxin up-regulated RNA* (*SAUR*) genes. Overall, the upregulation of auxin-related genes suggests that auxin signaling plays a role in regulating domatia development.

### Amino acid and carbohydrate transport and carbohydrate metabolism are upregulated in domatia

We observed multiple genes involved with amino acid transport, carbohydrate transport, and carbohydrate metabolism upregulated in domatia in both genotypes. Both genotypes were enriched for amino acid transport (GO:0006865) and amino acid transmembrane transport (GO:0003333), and several amino acid transporters are upregulated in domatia (Fig. 6).

**Fig. 6.**
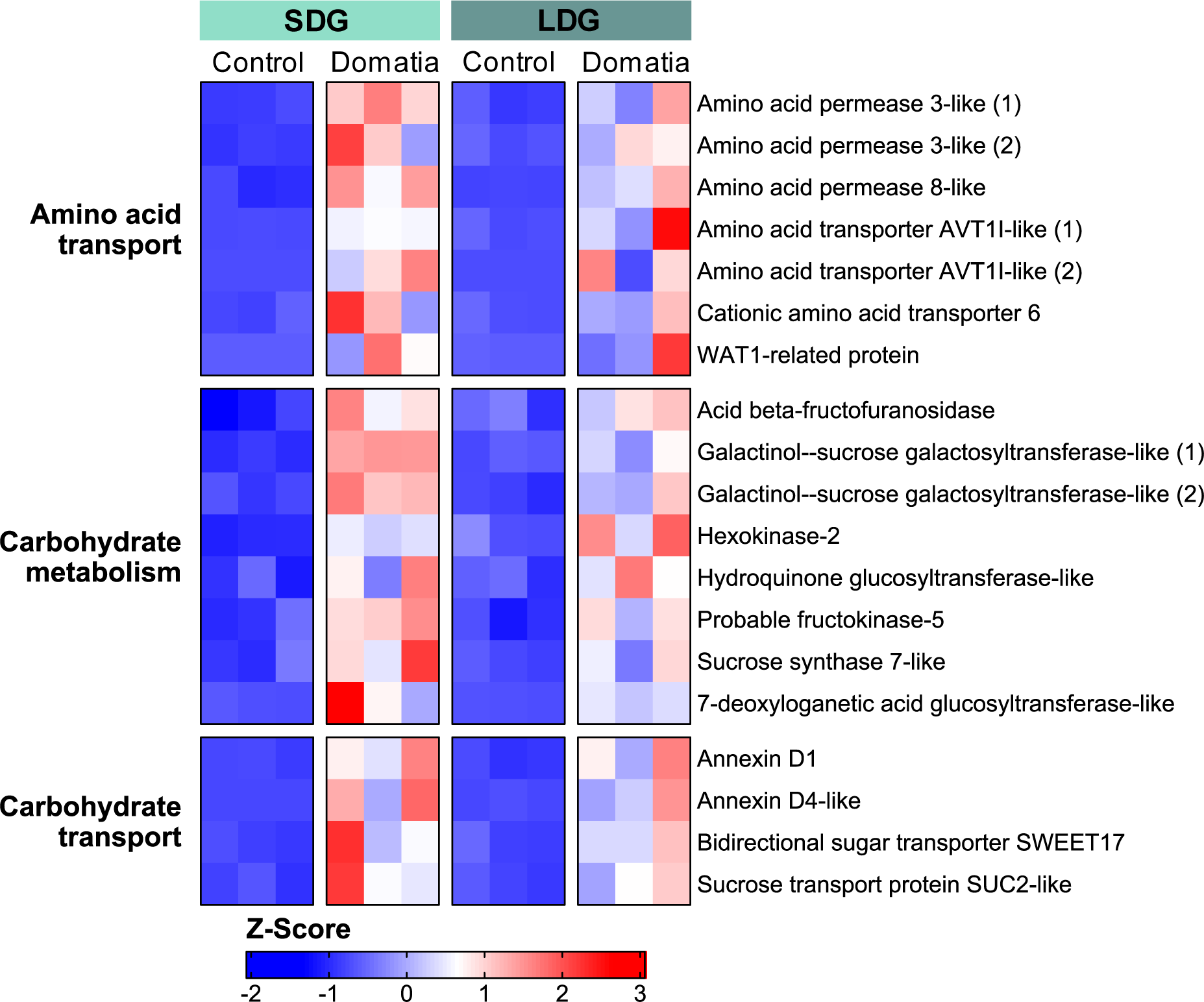
A heatmap of genes upregulated in domatia of both genotypes that are involved in the transport and metabolism of amino acids and carbohydrates. Each column represents a biological replicate. The cells are colored by Z-score, with blue representing lower expression and red representing higher expression.

Domatia also have higher expression of genes involved in carbohydrate metabolism and transport. While many of these genes overlap with those involved in hemicellulose biosynthesis, genes explicitly involved in general sugar metabolism and transport are also upregulated. We see evidence for the metabolism of starch, sucrose, and hexoses through the upregulation of the genes encoding products such as sucrose synthase 7-like, hexokinase-2, and fructokinase-5. Genes involved in the production of secondary metabolites specifically are upregulated in domatia, including two genes encoding galactinol--sucrose galactosyltransferase-like, orthologous to the raffinose synthase AtRS5. Additionally, genes encoding sugar transporters, such as SWEET17, a SUC2-like protein, and Annexin D1, are upregulated (Fig. 6). Several of these genes are closely related to genes upregulated in the extrafloral nectaries (EFNs) of cotton (*Gossypium hirsutum*) (Chatt *et al*., 2021). Like domatia, EFNs are mutualistic structures produced by plants, but EFNs provide nectar (rather than housing) to arthropods in return for protection and so the reason for this overlap is unclear. As EFNs are homologous to floral nectaries in many angiosperms (Lee *et al*., 2005; Weber & Keeler, 2013), we hypothesized that domatia-upregulated genes could also be expressed in grapevine floral tissues and involved in floral development. Supporting this, upregulated genes in LDG domatia are enriched for flower morphogenesis (GO:0048439) (Supporting Information Table S7). Comparing the expression of *V. vinifera* orthologs to domatia-upregulated gene expression in floral, leaf, and stem tissue revealed that 98.6% of orthologs were expressed in floral tissue and identified 17 *V. vinifera* orthologs that demonstrated strong preferential expression in floral tissue compared to both leaf and stem tissue (Supporting Information Methods S1 and Fig. S7). Interestingly, one of the orthologs identified was the transcription factor *VviAGL6a*, which has been shown to play a role in grapevine floral development (Palumbo *et al*., 2019) and grapevine gall development from phylloxera infection (Schultz *et al*., 2019).

### Intraspecific variation in domatia development

To understand intraspecific variation in domatia size, we identified genes where differences in expression levels between control and domatia tissue differed between the two genotypes. Nineteen genes showed significant genotype-tissue interactions (Fig. 7).

**Fig. 7.**
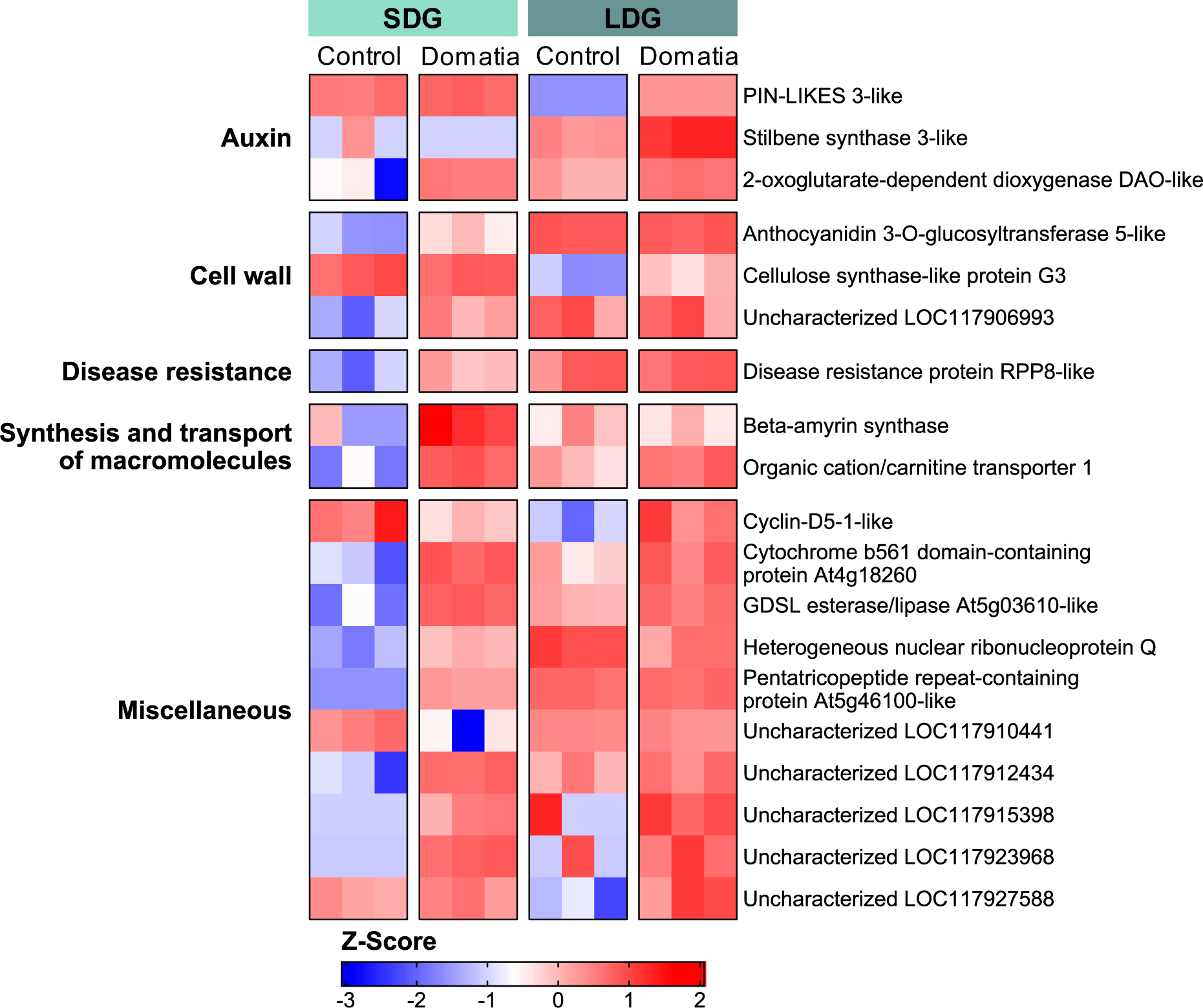
A heatmap of genes with significant domatia-genotype interactions between SDG and LDG. Each column represents a biological replicate. The cells are colored by Z-score, with blue representing lower expression and red representing higher expression.

Of these genes, 14 exhibited tissue-specific expression in SDG only, and five genes exhibited the opposite pattern with tissue-specific expression in LDG only. Nine of these genes are of unknown function, including LOC117912434 and LOC117927588, which are both orthologous to the Arabidopsis gene AT3G18670 and exhibit opposing patterns (the former being “domatia-responsive” in SDG and the latter in LDG) suggesting they may be functionally similar in the two genotypes.

Aside from pathways already implicated in domatia development, cell cycle regulation is implicated in intraspecific variation in domatia development due to the expression of the gene encoding Cyclin-D5-1-like, demonstrating genotype-tissue interactions. The Arabidopsis ortholog for this gene, *AtCYCD5;1,* is part of the D-type cyclin family implicated in regulating DNA replication, the cell cycle, and cellular differentiation (Schnittger *et al*., 2002; Dewitte *et al*., 2003).

The other ten genes that exhibited significant genotype-tissue interactions were involved in the processes mentioned above. The gene encoding 2-oxoglutarate-dependent dioxygenase DAO-like, which is orthologous to the gene *AtDAO1* involved in auxin oxidation and auxin-JA crosstalk (Lakehal *et al*., 2019), exhibited tissue-specific expression in SDG only. The gene encoding Anthocyanidin 3-O-glucosyltransferase 5-like, likely involved in the biosynthesis of the precursors of lignin, and the uncharacterized gene LOC117906993, orthologous to *AtWAK2*, both exhibit tissue-specific expression in SDG only. However, two auxin transporters, the genes encoding stilbene synthase 3-like and PIN-LIKES 3-like, exhibited tissue-specific expression in LDG only. The gene encoding a cellulose synthase-like protein G3, likely involved in hemicellulose synthesis, exhibited tissue-specific expression in LDG only. Genes exhibiting genotype-tissue interactions involved in disease resistance and the synthesis or transport of molecules only exhibit tissue-specific expression in SDG.

Despite only 19 genes being differentially expressed between SDG domatia and LDG domatia, the two domatia genotypes differ greatly in both phenotype and transcripts upregulated in domatia (when compared to the control tissue). While there is overlap in the genes upregulated in domatia, a considerable number of genes (909 in SDG domatia and 221 in LDG domatia) are not shared when comparing expression between domatia and control tissue between the two genotypes. Of genes differentially expressed in domatia, 35-39% (459 genes in SDG and 227 in LDG) are differentially expressed between the control tissue of the two genotypes (Supporting Information Fig. S3). Most of these genes (80.6-86.2%) exhibit a specific pattern where they: a) show lower expression levels in the control tissue of one genotype compared to the control leaf tissue of another genotype but are upregulated in domatia tissue of that particular genotype (in contrast to control tissue), while b) the other genotype demonstrates no difference in gene expression levels between control and domatia tissue. This suggests that differences in leaf tissue development may drive differences in domatia traits between genotypes of *V. riparia*. Genes that match this pattern in SDG domatia tissue are enriched for meristem and tissue development (*P* < 0.05), which supports this as well (Supporting Information Fig. S8).

To better understand the relationship between domatia traits and leaf development, we landmarked leaves from SDG and LDG plants to see if they differed in overall leaf shape, which would demonstrate differences in leaf development between the two genotypes. Landmarking leaves from SDG and LDG plants revealed that the two genotypes are significantly different in leaf shape (H = 2.96, *P* = 0.009), with SDG leaves having narrower lateral and apical lobes and a narrower upper lateral sinus than LDG (Fig. 8 and Supporting Information Fig. S9).

**Fig. 8.**
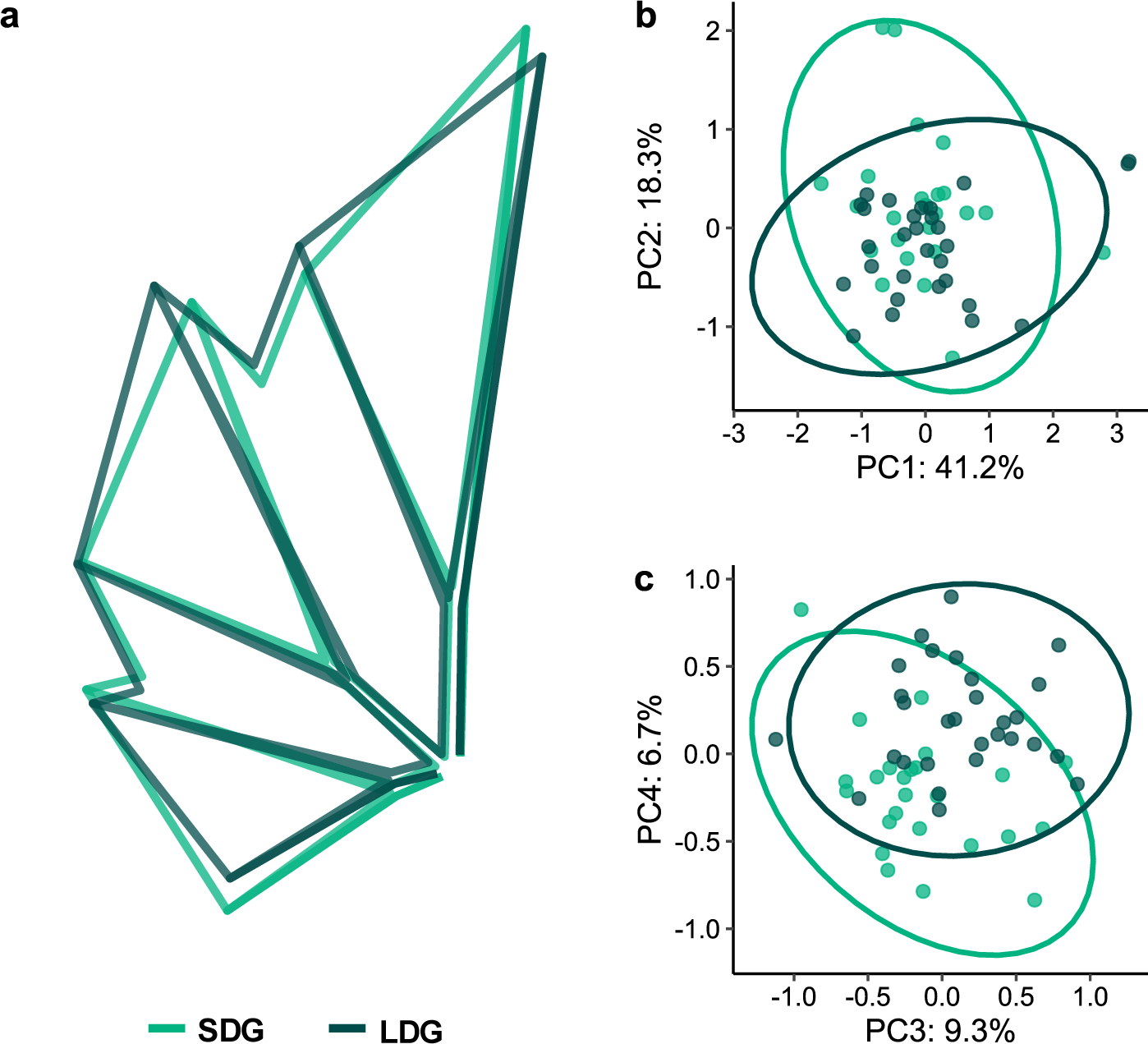
Differences in leaf shape between SDG and LDG. **(a)** Mean leaf shapes of SDG leaves (light green) and LDG leaves (dark green) rotated and scaled identically. **(b and c)** Principal component analysis (PCA) of leaf shapes, SDG in light green and LDG in dark green. **(b)** PCs 1 and 2, with PC1 contributing to 41.2% of variation and PC2 to 18.3%. **(c)** PCs 3 and 4, with PC3 contributing to 9.3% of variation and PC4 to 6.7%. In Fig. S9, Eigenleaves display the morphological characteristics of each PC.

These differences in leaf shape reflect altered angles of vein axils on the leaves where domatia form, which could indirectly impact domatia development and influence domatia size. It is also possible that the genetic differences between genotypes drive differences in leaf shape and are directly responsible for differences in domatia size.

## DISCUSSION

Understanding the genetic underpinnings of ecologically important traits is a central goal linking subfields in biology, yet the genetic bases of many ecologically relevant traits remain understudied. Here, we present the first transcriptomic study aimed at understanding the genetic drivers of the development of mite domatia, small structures on the undersides of plant leaves that mediate a powerful and pervasive mutualism between plants and beneficial mites. Several of the genes we identified overlap with genes previously implicated in domatia development in *Vitis,* including *V. riparia* genes encoding TFs thought to regulate trichome development (Barba *et al*., 2019; LaPlante *et al*., 2021), as well as Importin Alpha Isoform 1 and GATA Transcription Factor 8 (LaPlante *et al*., 2021). We found that genes related to domatia development are similar to those involved in trichome and tuber domatia development, including genes involved in trichome regulation, cell wall biosynthesis/modification, plant hormone signaling, and biotic responses. We also found that amino acid transport and carbohydrate metabolism/transport are implicated in the development of domatia. These findings provide insight into the genetic drivers and functioning of this important phenotype.

### C2H2 ZFPs and SPLs are likely key regulators of trichome initiation in *Vitis* domatia

Trichomes have convergently evolved in numerous angiosperm lineages and are regulated by distinct pathways between species (Serna & Martin, 2006; Han *et al*., 2022), and the genetic pathways of trichome development in *V. riparia* have yet to be uncovered. Our findings, previous genetic research in *Vitis* (Barba *et al*., 2019; LaPlante *et al*., 2021), and studies of trichome development in other species (Han *et al*., 2022), all support C2H2 ZFPs and SPLs as probable key trichome regulators in *V. riparia* and related species. One of the C2H2 ZFPs identified in previous studies in *Vitis* is orthologous to *AtZFP6*, whose *V. riparia* ortholog was upregulated in domatia in both genotypes in this study. This *V. vinifera* gene, VIT_205s0077g01390, was identified as a candidate gene for hair on leaf blades (Barba *et al*., 2019) and domatia density (LaPlante *et al*., 2021). Two additional ZFPs were identified as genetic candidates for leaf trichome traits by Barba *et al*. (2019). Two *SPLs* were also upregulated in the domatia we sequenced, and Barba *et al*. (2019) linked the *V. vinifera* gene VIT_15s0021g02290, orthologous to *AtSPL8* in Arabidopsis, to domatia size. The overlap between C2H2 ZFPs and SPLs in our dataset, as well as previous quantitative genetic studies in *Vitis* (Barba *et al*., 2019; LaPlante *et al*., 2021), suggest that these TFs may play an essential role in regulating trichome and/or domatia development in *Vitis*.

### Insights into *Vitis* domatia cell wall biosynthesis and composition

Our findings provided insight into the biosynthesis and composition of cell walls in domatia. While our domatia samples include both laminar tissue and trichomes, we expect trichomes to have vastly different cell wall composition compared to laminar tissue. Accordingly, DEGs in domatia involved in cell wall biosynthesis provide insight into the composition of trichome cell walls. Gene pathways involved in xyloglucan, xylan, pectin, and lignin biosynthesis were upregulated in domatia tissue. While xyloglucan, pectin, and lignin are fairly common components of cell walls in both normal leaf tissue and trichomes (Marks *et al*., 2008; Bowling *et al*., 2011), xylan is not (Bowling *et al*., 2008, 2011). However, xylan is important for general protection against herbivores and pathogens (Gao *et al*., 2017; Joo *et al*., 2021). A previous study investigating loci associated with downy mildew resistance in grapevine also identified a candidate gene involved in xylan biosynthesis (Divilov *et al*., 2018). It is possible that xylan production in domatia trichomes in *Vitis* enables protection against downy mildew either directly or indirectly through facilitating mite mutualisms. It is also possible that the upregulation of xylan-related genes is due to the closeness of vascular tissue to domatia and trichomes (Tilney *et al*., 2012; Gago *et al*., 2016), which typically have large amounts of xylan (Moore *et al*., 2014). Future work characterizing the composition of cell walls in domatia trichomes would clarify the specific role of these genes in domatia.

### Auxin and JA mediate the development of domatia in *V. riparia*

We found that auxin and JA genes are heavily upregulated during domatia development. The upregulation of auxin genes is unsurprising, as auxin plays significant roles in cell elongation, leaf trichome development (Xuan *et al*., 2020) and tuber domatium development (Pu *et al*., 2021). However, to our knowledge, a connection between JA and domatia has not been studied or shown. Notably, JA is implicated in other plant-arthropod mutualisms, increasing nectar secretion in EFNs (Heil *et al*., 2001, 2004; Kost & Heil, 2008; Hernandez-Cumplido *et al*., 2016). JA may mediate plant-mite interactions in some cases, similar to how JA mediates ant-plant interactions in some EFN-bearing species (Heil *et al*., 2001, 2004; Kost & Heil, 2008; Hernandez-Cumplido *et al*., 2016). JA could induce the structural development of domatia by regulating trichome development, as shown in a few other species (Han *et al*., 2022). It could also regulate the release of plant volatiles, as has been shown in other systems (Schmelz *et al*., 2003; Ament *et al*., 2004; Degenhardt *et al*., 2010), and these volatiles could mediate mutualisms with mites through signaling or provide another layer of direct defense (Baldwin, 2010). Alternatively, JA could provide direct defense against bacteria and fungi growing on mite waste within domatia (excrement, exoskeletons, etc.). Future work investigating the impact of JA application on *V. riparia* could clarify JA’s role in domatia.

### Insights into mite-plant mutualisms in *Vitis*

Our findings provide insight into possible ways *V. riparia* domatia mediate mite mutualisms. We saw evidence for volatile production through the expression of genes involved in terpenoid synthesis. This includes the ortholog to TERPENE SYNTHASE 21 (AtTPS21) in Arabidopsis. AtTPS21 is involved in the production of (*E*)-β-caryophyllene (Chen *et al*., 2003a), which mediates both direct defense against microbial pathogens (Cowan, 1999; Huang *et al*., 2012) and indirect defense against herbivores by attracting natural enemies (Rasmann *et al*., 2005; Köllner *et al*., 2008). It is possible that this volatile attracts mites to domatia or is a direct defense to inhibit pathogen growth within domatia.

We also see evidence for methyl salicylate (MeSa) emission through the upregulation of two genes encoding salicylate carboxymethyltransferase-like and one encoding 7-methylxanthosine synthase 1-like, all of which are orthologous to *AtBSMT1* which is responsible for MeSa production (Chen *et al*., 2003b). MeSa is a common plant volatile typically released after herbivory (Chen *et al*., 2003b; Snoeren *et al*., 2010) that repels herbivores (Koschier *et al*., 2007; Ulland *et al*., 2008) and attracts predators (De Boer & Dicke, 2004; James & Price, 2004; Mallinger *et al*., 2011). In grapevine, MeSa attracts the predaceous mite *Typhlodromus pyri* (Gadino *et al*., 2012) that inhabits leaves (English-Loeb *et al*., 2002). The upregulation of the *V. riparia* orthologs to *AtBSMT1* suggests that MeSa production and emission may occur in domatia, which could attract predatory mites. Due to their small size, it is challenging to capture domatia-specific volatiles. However, future work investigating volatile emissions from domatia could test hypotheses surrounding the mediation of domatia inhabitancy through volatiles.

### Gene expression patterns in domatia share similarities with EFNs

Several genes we identified involved in macromolecule biosynthesis and transport are closely related to genes involved in development and nectar production in EFNs (Roy *et al*., 2017; Chatt *et al*., 2021). We found that upregulated genes in LDG domatia are enriched for flower morphogenesis, and *V. vinifera* orthologs of genes upregulated in domatia were expressed in floral tissue compared to leaf and stem tissue. Further, one of the orthologs identified was also related to grapevine gall development from phylloxera infection (Schultz *et al*., 2019), suggesting that floral pathways have been co-opted in different ways to enable the development of diverse structures like EFNs, galls, and domatia. Understanding the overlap of domatia genes with genes involved in EFN, gall, and floral development may provide insight into potential pathways modified to enable the evolution of plant structures that mediate mutualisms.

The overlap between genes upregulated in domatia and EFNs could also be due to functional similarities. To our knowledge, no studies have suggested that *V. riparia* domatia produce secretions for beneficial mites. However, nectar applied to *V. riparia* leaves increased mite recruitment (Weber *et al*., 2016), and there is evidence of material exchange between mites and domatia in *Plectroniella armata* (Tilney *et al*., 2012). The considerable upregulation of sugar and amino acid transport genes (Fig. 6) and the upregulation of many genes involved in floral development (Supporting Information Fig. S7) suggests the possibility that these phenomena could be due to material exchange from the plants to the mites and begs future studies investigating the detailed morphologies and functions of *V. riparia* domatia. Alternatively, like EFNs (Chatt *et al*., 2021), domatia and grapevine trichomes tend to be located near vascular tissue bundles (Tilney *et al*., 2012; Gago *et al*., 2016), so macromolecule biosynthesis and transport gene upregulation may be due to an abundance of vascular tissue in domatia samples.

### Intraspecific variation in domatia size may be due to differences in leaf development

Despite varying substantially in domatia size (Fig. 1c), with LDG domatia nearly two times larger than SDG domatia, we only identified 19 genes differentially expressed between SDG and LDG domatia. However, many genes involved in domatia development overlap with genes differentially expressed between SDG and LDG control tissue (35-39%) (Supporting Information Fig. S3). Further, the two genotypes varied in domatia traits and overall leaf shape (Fig. 8 and Supporting Information Fig. S9). Thus, differences in overall leaf development may shape domatia traits in *V. riparia*. Previous studies in *Vitis* and *Begonia* found a relationship between leaf morphology and leaf trichomes (McLellan, 2005; Chitwood *et al*., 2014), and Barba *et al*. (2019) also suggest a link between leaf morphology and trichome development in *Vitis*. Future work investigating domatia and leaf development in *Vitis* together could unravel the molecular genetic mechanisms and developmental processes enabling the potential link between leaf morphology and domatia.

## ACKNOWLEDGEMENTS

We would like to thank Grace Fleming, Bruce Martin, Erika LaPlante, Andrew Myers, and other members of the Weber lab for helpful conversations surrounding these research ideas. We are grateful to Dan Chitwood, Emily Josephs, and Robin Buell for helpful discussions on this work and feedback on this manuscript. We thank David Lowry and his research group for feedback on this manuscript as well. We are grateful to the Genomics Core at Michigan State University and the Institute for Cyber-Enabled Research at Michigan State University for their services. We are also grateful to the staff of the Research Greenhouse Complex at Michigan State University and the staff of the research greenhouses at the University of Michigan for their assistance with plant growth. This work was supported by NSF DEB-1831164 (awarded to MGW). EJR was supported by the University Distinguished Fellowship at Michigan State University.

## AUTHOR CONTRIBUTIONS

MW, CN, and EJR conceptualized and designed the study. CG provided plant material and characterized domatia traits. EJR carried out the RNA-seq analysis and performed leaf landmarking. MW, CN, and EJR assisted in data interpretation. EJR wrote the first draft of the manuscript. All authors assisted with final drafts of the manuscript.

## DATA AVAILABILITY

Leaf scans, domatia phenotype data, and raw gene expression counts from this study are openly available in Dryad at XXX, reference number XXX. RNA sequencing data from this study are provided on the NCBI Sequence Read Archive under BioProject PRJNA1083535. The code used for data analysis in this study is available on GitHub: https://github.com/eleanore-ritter/domatia-transcriptome.

## SUPPORTING INFORMATION

**Fig. S1** Domatia size during leaf expansion

**Fig. S2** Sampling schematic for tissue collected for RNA-sequencing

**Fig. S3** The number of differentially expressed that overlap between comparisons

**Fig. S4** Cellular Component GO terms enriched in genes upregulated in domatia

**Fig. S5** Molecular Function GO terms enriched in genes upregulated in domatia

**Fig. S6** C2H2 ZFPs and SPLs upregulated in developing domatia

**Fig. S7** Genes upregulated in domatia expressed primarily in floral tissue

**Fig. S8** Biological Process GO terms enriched in domatia-responsive genes from SDG that are not domatia-responsive in LDG

**Fig. S9** Eigenleaves from the PCA comparing leaf shape between scaled SDG and LDG leaves, for PC 1-4

**Methods S1** Methods for comparing the expression of V. vinifera orthologs of genes involved in domatia development across various tissue types

**Table S1** Read counts and mapping rates for all RNA-sequencing samples

**Table S2** Differentially expressed genes between SDG control tissue and LDG control tissue

**Table S3** Differentially expressed genes between SDG control tissue and SDG domatia tissue

**Table S4** Differentially expressed genes between LDG control tissue and LDG domatia tissue

**Table S5** Genes that demonstrated significant domatia-genotype interactions between SDG and LDG

**Table S6** Biological Process GO terms enriched in genes upregulated in SDG domatia

**Table S7** Biological Process GO terms enriched in genes upregulated in LDG domatia

