## Supplemental Figures and Methods for "Small, but mitey: Investigating the molecular genetic basis for mite domatia development and intraspecific variation in *Vitis riparia* using transcriptomics"

The following Supporting Information is available for this article:

**Fig. S1** Domatia size during leaf expansion. Leaf width (cm) and domatium radius (mm) of leaves collected from bud burst to full expansion for both genotypes of *V. riparia*, 588710 (SDG) and 588711 (LDG). The blue vertical lines represent the lower and upper cutoffs for the leaves that domatia and control tissue were collected from for RNA-sequencing.

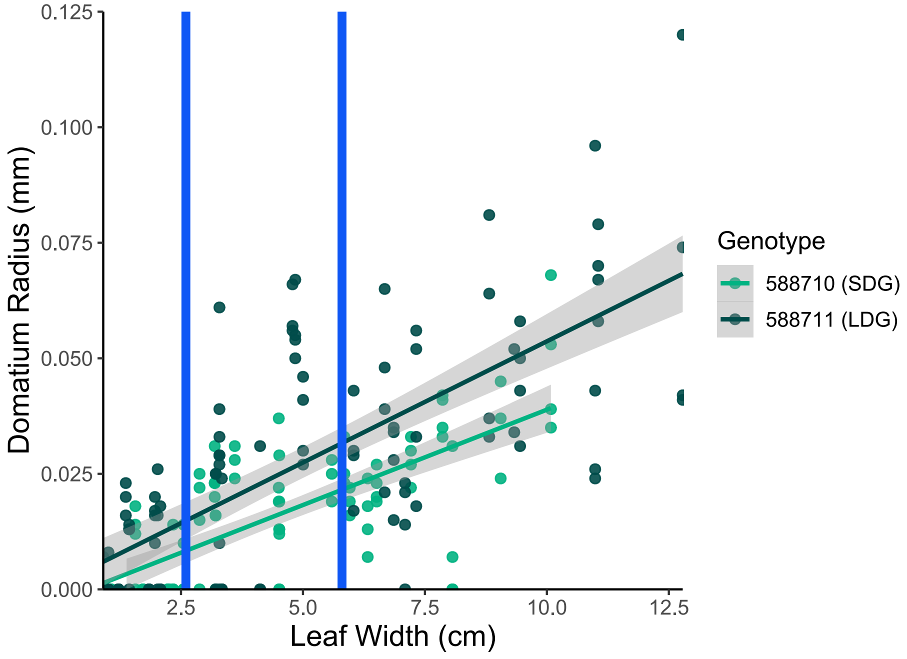


**Fig. S2** Sampling schematic for tissue collected for RNA-sequencing. Domatia and control tissue were collected using a 1.5 mm hole puncher to take circular tissue samples of leaves. The domatia samples were collected by punching out the domatia, avoiding the veins nearby, while the control samples were collected by punching out tissue 1.5 mm directly out from the domatia. The pink circle demonstrates where domatia tissue were collected, and the black circle demonstrates where control tissue was collected.

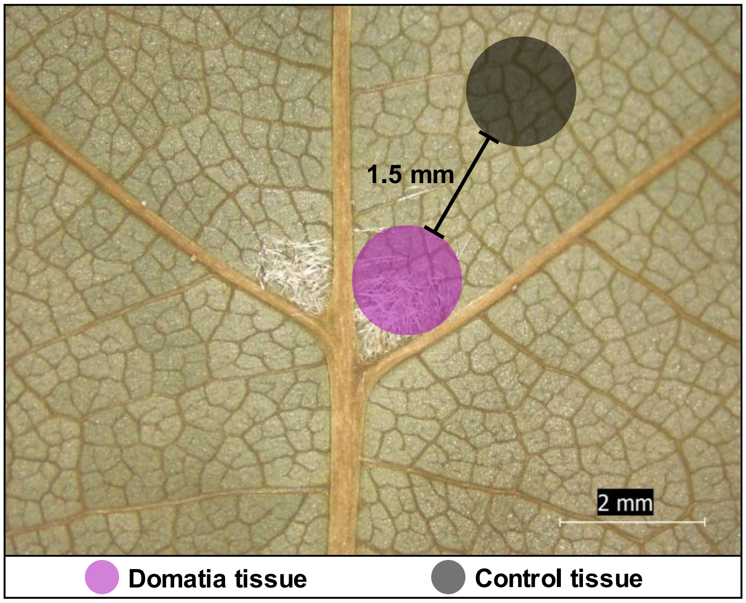


**Fig. S3** The number of differentially expressed that overlap between comparisons. From left to right: control SDG tissue vs domatia SDG tissue (SDG: C vs D), control SDG tissue vs control LDG tissue (Control vs Control), domatia SDG tissue vs domatia LDG tissue (Domatia vs Domatia), and control LDG tissue vs domatia LDG tissue (LDG: C vs D).

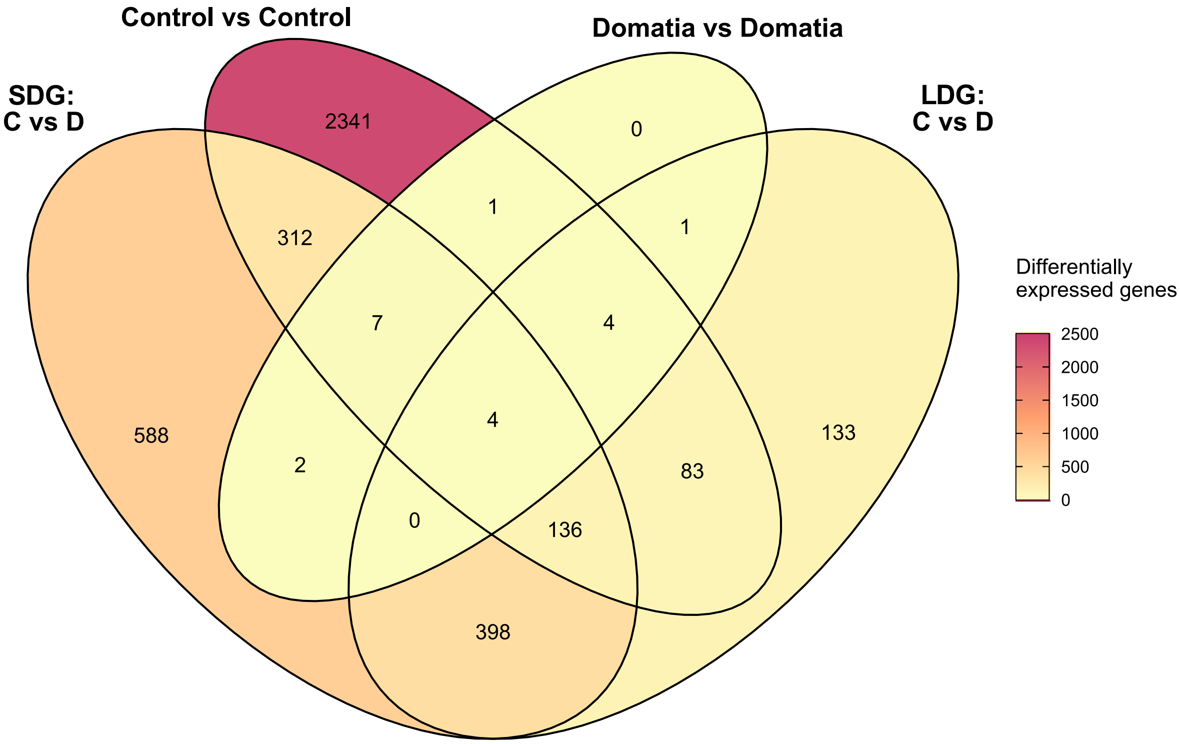

**Fig. S4** Cellular Component GO terms enriched in genes upregulated in domatia. Cellular Component gene ontology (GO) terms enriched ($P < 0.05$) in genes differentially expressed between domatia tissue and control tissue that overlap between both genotypes.

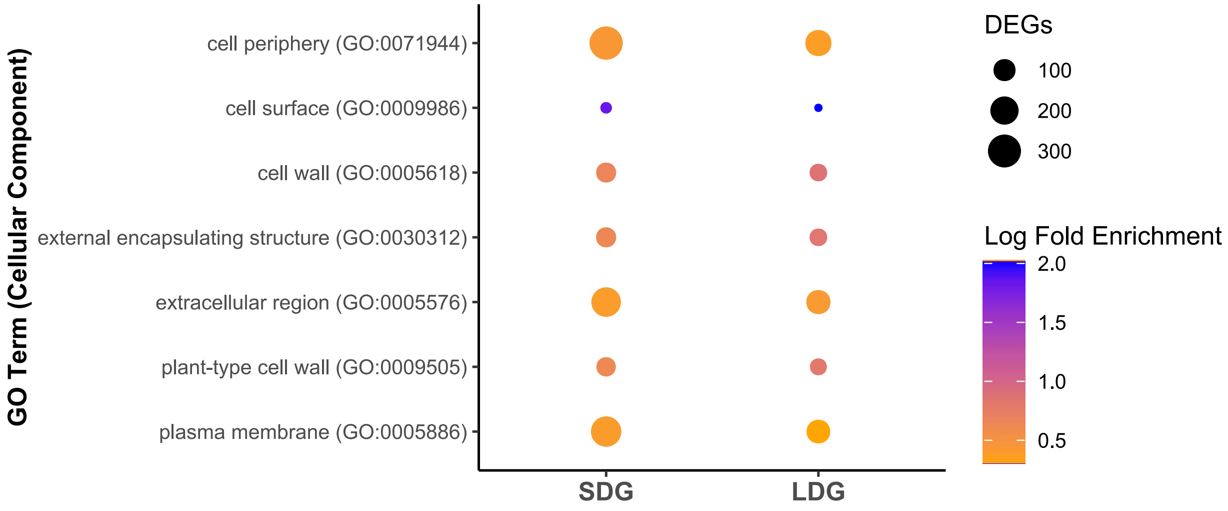


**Fig. S5** Molecular Function GO terms enriched in genes upregulated in domatia. Molecular Function gene ontology (GO) terms enriched ($P < 0.05$) in genes differentially expressed between domatia tissue and control tissue that overlap between both genotypes.

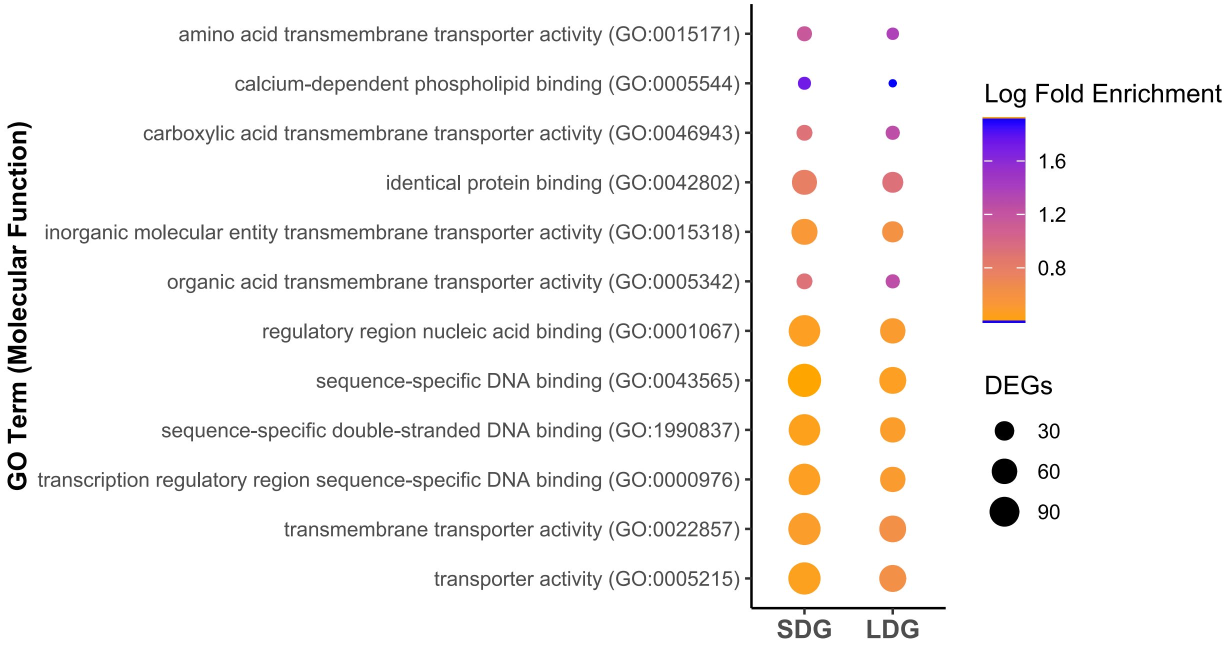

**Fig. S6** C2H2 ZFPs and SPLs upregulated in developing domatia. A heatmap of C2H2 ZFPs and SPLs upregulated in domatia of either both genotypes or one of the genotypes. Each column represents a biological replicate. The cells are colored by Z-score, with blue representing lower expression and red representing higher expression. Genes not differentially expressed in domatia of a particular genotype are denoted with an asterisk.

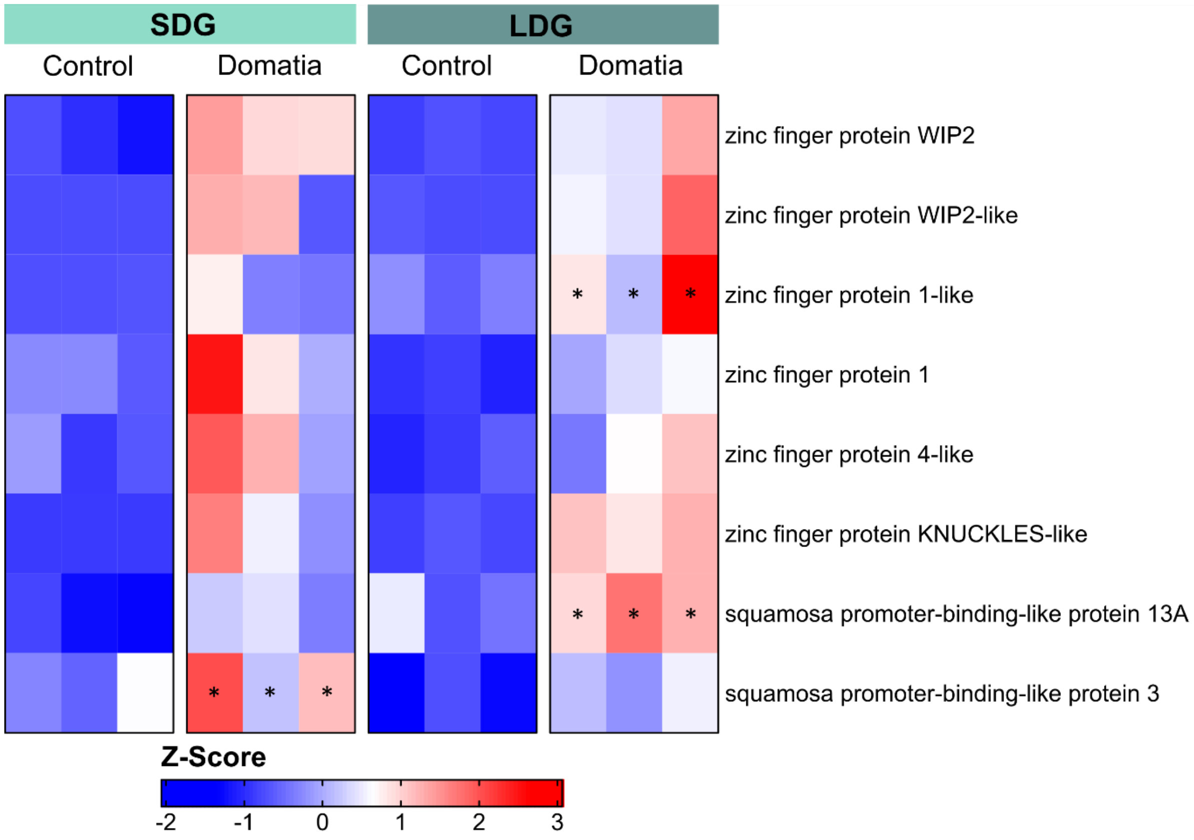


**Methods S1** Methods for comparing the expression of *V. vinifera* orthologs of genes involved in domatia development across various tissue types.

Because analyses revealed evidence of genes involved in floral structures in domatia, we asked whether genes involved in domatia development showed specific expression patterns in other tissue types in *Vitis*. We identified *V. vinifera* orthologs to genes differentially expressed in domatia using Diamond v0.8.36 (Buchfink *et al.*, 2015) and the *V. vinifera* reference genome 12X.v2 and its annotation VCost.v3 (Canaguier *et al.*, 2017). The expression patterns of these genes were compared in floral (inflorescence), leaf, and stem tissue using GREAT (GRape Expression ATlas) (https://great.colmar.inrae.fr/, unpublished). Genes were identified as being expressed in floral tissue when they had more than 17.45 reads (the 25% quantile of read counts in dataset) expressed in a floral tissue sample. The expression of genes displaying preferential expression in floral tissue was normalized using a Z-score.

**Fig. S7** Genes upregulated in domatia expressed primarily in floral tissue. A heatmap of the expression of *V. vinifera* orthologs to domatia-upregulated genes for expression that demonstrate preferential expression in floral tissue compared to leaf and stem tissue. Each column represents a biological replicate from GREAT (GRape Expression ATlas). The colors in each cell represent the Z-score, with white representing lower expression and red representing higher expression.

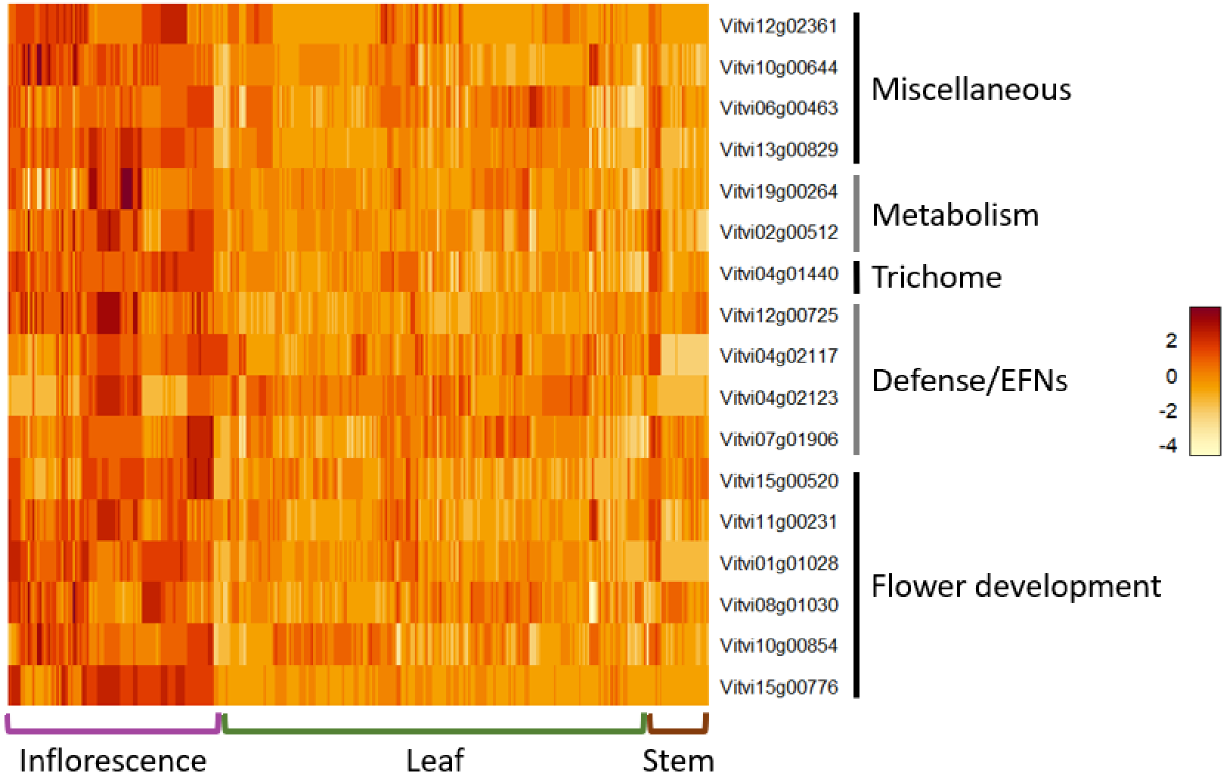


**Fig. S8** Biological Process GO terms enriched in domatia-responsive genes from SDG that are not domatia-responsive in LDG. Biological Process gene ontology (GO) terms enriched ($P < 0.05$) in genes differentially expressed only in SDG domatia that match the following pattern: a) show lower expression levels in the control tissue of SDG compared to the control leaf tissue of LDG, and are upregulated in domatia tissue of SDG (in contrast to control tissue), while b) LDG demonstrates no difference in gene expression levels between control and domatia tissue.

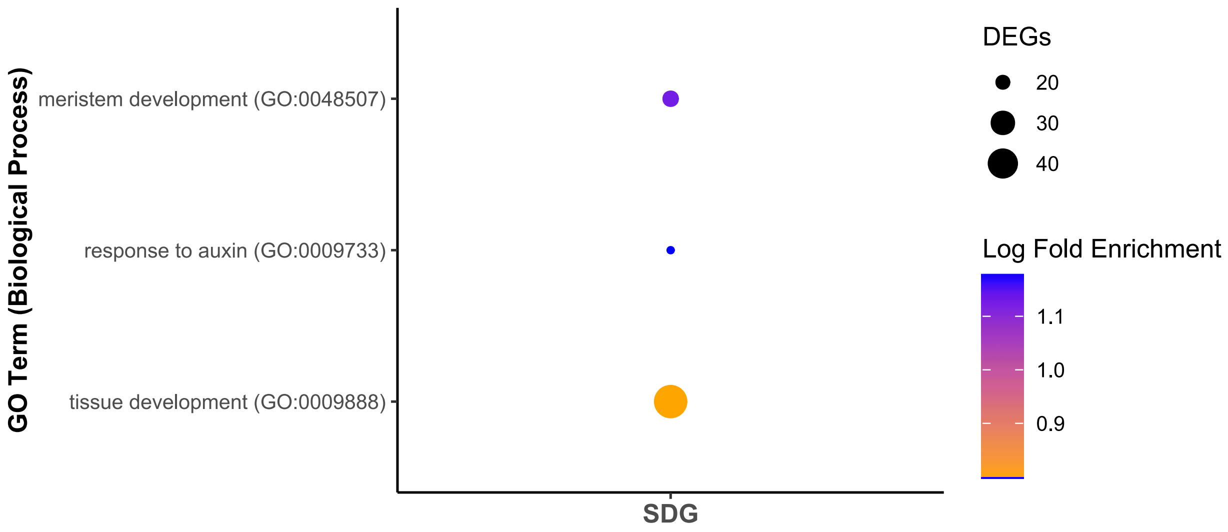


**Fig. S9** Eigenleaves from the PCA comparing leaf shape between scaled SDG and LDG leaves, for PC 1-4.

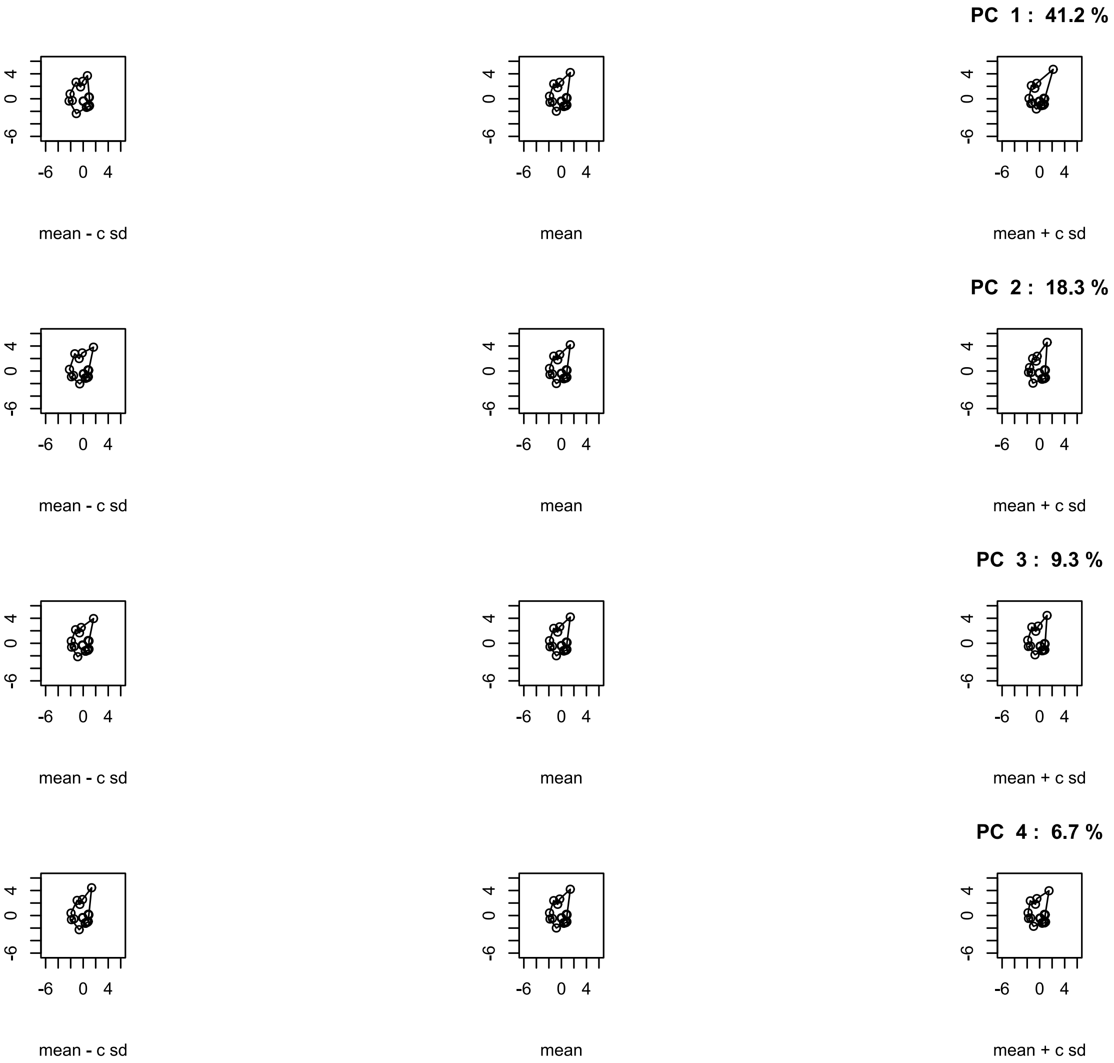
PC 1 : 41.2 %
mean - c sd
mean
mean + c sd
PC 2 : 18.3 %
mean - c sd
mean
mean + c sd
PC 3 : 9.3 %
mean - c sd
mean
mean + c sd
PC 4 : 6.7 %
mean - c sd
mean
mean + c sd
